# A surface-intrinsic framework for topology-preserving hippocampal alignment and precision morphometry

**DOI:** 10.64898/2026.07.22.740127

**Authors:** J DeKraker, D Bansal, M Snyder, BG Karat, N Talaei, M Salman, A Ngo, J Chen, E Sahlas, J Royer, DG Cabalo, MF Glasser, T Coalson, J Harwell, M Torkamani-Azar, Y Liu, J Tohka, JC Lau, AC Evans, BC Bernhardt, AR Khan

**Author notes:** co-shared authors (JC Lau, AC Evans, BC Bernhardt, AR Khan).

## Abstract

Accurate alignment of hippocampal anatomy across individuals remains challenging due to complex and highly variable folding patterns that are not well captured by conventional volumetric approaches. HippUnfold introduced a surface-based representation of the hippocampus, but key components—including coordinate estimation and inter-subject correspondence—were defined in the volumetric domain, making them susceptible to topological errors and interpolation artifacts. Here, we introduce a surface-intrinsic formulation of hippocampal unfolding in which geometry, intrinsic coordinates, and correspondence are defined directly on subject-specific surface manifolds. Intrinsic anterior–posterior and proximal–distal coordinates are computed by solving Laplace equations on the surface, and correspondence is established through surface-based resampling in unfolded space, replacing inverse volumetric warping. Relative to the original HippUnfold approach, this formulation improves test–retest consistency, subject identifiability, and mesh quality, while better preserving subject-specific gyral and sulcal morphology. Surface representations show reduced distortion between folded and unfolded spaces and eliminate misplaced or outlier vertices associated with volumetric warping. These improvements translate to enhanced sensitivity in a clinical application, improving lateralization of temporal lobe epilepsy. These results demonstrate that a surface-intrinsic formulation provides a principled and robust foundation for hippocampal unfolding, enabling topology-preserving alignment and more accurate characterization of inter-individual variability in health and disease.

## Introduction

The hippocampus is an archicortical precursor to the neocortex and is preserved across all mammals, with homologues or analogues in most species exhibiting experience-dependent behavioural flexibility (Rubin et al. 2014). It has important roles in memory, spatial navigation, and prediction, and may be a foundational structure informing neocortical learning (Stachenfeld et al. 2017; DeKraker et al. 2025). Despite being one of the most heavily studied brain structures, accurately modeling hippocampal shape and aligning hippocampi across individuals remains a major challenge. This difficulty arises from the complex and highly variable folding patterns of the hippocampus, which, like the neocortex, exhibits substantial inter-individual variability in gyral and sulcal morphology. For a time this was underappreciated in human studies based on coronal sections, in both health and disease (Ding and Van Hoesen 2015; Blümcke et al. 2013), but in more recent work, 3D analyses reveal that gyrified folding patterns extend throughout the hippocampus in many individuals (DeKraker et al. 2020, 2023, 2025; Cai et al. 2019). In the neocortex, such variability is addressed using surface-based methods that align individuals in a flattened or inflated space, avoiding collisions due to differences in folding patterns (Van Essen et al. 1998; DeKraker et al. 2021; Robinson et al. 2014). Similar approaches have been proposed for the hippocampus (Van Essen et al. 1998; DeKraker et al. 2021; Robinson et al. 2014; Kim et al. 2014), highlighting the importance of topology-aware representations for cross-subject alignment.

HippUnfold (DeKraker et al. 2022) is an automated framework for the segmentation and surface-based unfolding of the hippocampus in MRI, designed to address the variability in hippocampal folding across individuals. It works by indexing the hippocampus along its “natural” anterior-posterior and proximal-distal axes (DeKraker et al. 2018; Paquola et al. 2020), enabling representation of hippocampal anatomy in a two-dimensional unfolded coordinate space. This unfolded space is then registered between individuals in 2D (DeKraker et al. 2023), similar to spherical registration often used in the neocortex (Klein et al. 2010; Fischl et al. 1999; Yeo et al. 2010). Each of these transforms is reversible, meaning that subjects can be aligned to one another in their native space or data can be represented on a flat or a highly smoothed “canonical” surface. By explicitly modeling hippocampal topology, HippUnfold overcomes key limitations of volumetric registration approaches, which are ill-suited to capturing inter-individual variability in gyral and sulcal patterning (Wisse et al. 2016; Hickling et al. 2024; Iglesias et al. 2015; Caldairou et al. 2016). However, despite adopting a surface-based representation, core components of the original HippUnfold framework, including coordinate estimation and mapping between folded and unfolded spaces, were initially derived from volumetric formulations. As a result, these steps remain susceptible to topological shortcuts, interpolation artifacts, and distortions that can impact vertex correspondence and surface geometry, particularly in regions of fine-scale folding.

Capitalizing on vertex-equivalence, we recently released HippoMaps, an open data warehouse that aggregates normative hippocampal data across imaging modalities, including *in vivo* and *ex vivo* MRI, histology, and functional imaging (DeKraker et al. 2025). This work further highlights the value of surface-based representations for integrating data across individuals with differing folding patterns and across imaging modalities. Together, these developments underscore the importance of topology-aware frameworks for hippocampal analysis, while motivating methods in which geometry, coordinates, and correspondence are defined intrinsically on the surface manifold.

Here, we reformulate hippocampal unfolding as a surface-intrinsic problem in which geometry, intrinsic coordinates, and inter-subject correspondence are defined directly on the hippocampal surface manifold. In contrast to prior implementations, where surfaces were derived from volumetric coordinate fields (DeKraker et al. 2022), intrinsic anterior–posterior and proximal–distal coordinates are computed by solving Laplace equations directly on subject-specific surface meshes, ensuring that coordinate systems respect hippocampal topology and avoid shortcuts across adjacent folds. Inter-subject correspondence is established through surface-based resampling in unfolded space, replacing inverse volumetric warping and preserving geometric fidelity without introducing interpolation artifacts or cross-depth smoothing. Together, these changes unify surface representation and computation within a single framework, resulting in improved preservation of subject-specific folding patterns, more consistent vertex correspondence, and reduced distortion between folded and unfolded spaces.

## Materials and Methods

### Overview of the hippocampal unfolding framework

The improved HippUnfold version consists of four main stages: 1) segmentation of hippocampal tissue types, 2) surface-fitting, 3) application of a 2D coordinate system (analogous to surface-based parameterization used in the neocortex), 4) surface-registration to a standard template in unfolded space. These steps are reviewed in **Figure 1**. The result is a set of subject-specific surfaces with vertex-wise correspondence across depths and between individuals, enabling consistent cross-subject alignment in both folded and unfolded representations. Vertices can then be parcellated according to a subfield atlas, continuous anterior-posterior gradient, or analyzed directly at the vertex-level.

**Figure 1.**
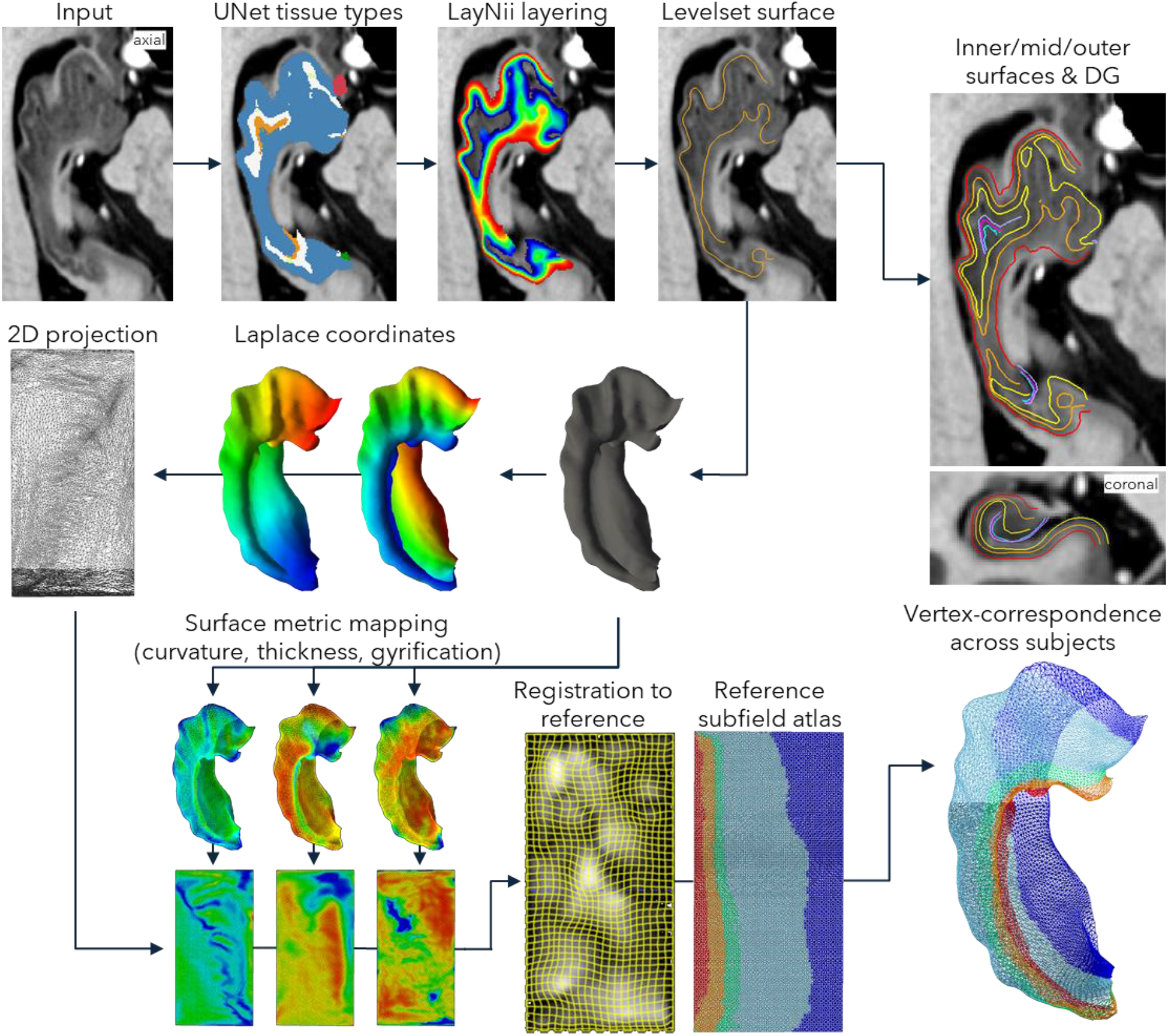
The HippUnfold version 2 pipeline. Input volumes are registered to a volumetric space (default CITI168 (Pauli et al. 2018)), enabling cropping, upsampling, and reorienting to coronal oblique to the hippocampal long axis around the left and right hippocampi. Cropped volumes are then fed into a nnUNet (Isensee et al. 2021) model trained for the given data type. This produces segmentation of tissue types including, critically, separation of folds via the strata radiatum and lacunosum moleculaire and background labels, as well as anterior termini on the hippocampal-amygdala transition area, posterior termini on the indiseum griseum, proximal termini on the neocortex, and distal termini on the dentate gyrus. Template shape injection (optional but enabled by default; not shown in the diagram) is used to post-process tissue segmentation. Layering is applied via LayNii (Huber et al. 2021) using the equidistance model. A levelset surface is generated at the 0.5 distance threshold. Edge vertices are then assigned to the anterior, posterior, proximal, and distal labels according to their Euclidean distance to the corresponding terminal labels, and these are used to solve two corresponding and approximately orthogonal Laplace fields, which serve as coordinates to map each vertex to a 2D space. In parallel, inner and outer surfaces are computed via translation of the midthickness surface across a warpfield between the inner (0-0.5) to outer (0.5-1) equidistance bins. From the midthickness, inner, and outer surfaces, morphometric measures of thickness, gyrification index, and curvature are computed. These features are then used to drive 2D unfolded registration to a single reference template with corresponding subfields (default multihist7 from previous work (DeKraker et al. 2023)). Resampling of the native midthickness, inner, and outer surfaces is then performed using standard surface tessellations (default 8k vertices), producing subject-specific surfaces with vertex-equivalence and interchangeable subfield atlases (default the maxprob from (DeKraker et al. 2023)). These surface operations are also performed in parallel on the dentate gyrus (without unfolded registration, meaning that vertex-equivalence is based solely on topological equivalence).

The following sections describe the methodological components underlying this framework, including surface construction, intrinsic coordinate estimation, and surface-based correspondence.

### Conceptual overview of surface-intrinsic formulation

In prior implementations, hippocampal unfolding was formulated in the volumetric domain, where intrinsic coordinate systems (anterior–posterior, proximal–distal, and inner–outer) were computed as Laplace fields spanning hippocampal gray matter. These fields defined a continuous mapping between native Cartesian coordinates and unfolded coordinate space, which was then used to transform a canonical surface into subject-specific native space via inverse interpolation. As a result, both coordinate estimation and inter-subject correspondence were implicitly defined through volumetric transformations.

In the present work, we reformulate hippocampal unfolding as a surface-intrinsic problem in which geometry, coordinates, and correspondence are defined directly on the hippocampal surface manifold. Rather than deriving surfaces from volumetric coordinate fields, we explicitly construct subject-specific surfaces and solve intrinsic coordinate systems on these meshes. Inter-subject correspondence is then established in unfolded space and mapped back to native space using surface-based resampling.

This shift decouples surface representation from volumetric transformations and ensures that all key operations—coordinate estimation, parameterization, and correspondence—are constrained to the intrinsic topology of the hippocampal surface. The following sections describe each component of this surface-intrinsic formulation in detail.

### Surface-intrinsic coordinate estimation via Laplace formulation

As described above, prior implementations derived intrinsic anterior–posterior and proximal–distal coordinates by solving Laplace equations in the volumetric domain. While this formulation produces smooth coordinate fields, it is susceptible to topological errors arising from voxel discretization and imperfections in tissue segmentation. In particular, when opposing banks of a sulcus are separated by distances comparable to or smaller than the voxel size, volumetric solutions can introduce unintended “short-circuits” or bridges across folds. These errors distort the coordinate system, leading to misplacement of surface vertices and subfield boundaries despite preserving overall hippocampal volume. Such distortions are often difficult to detect using conventional volumetric quality control metrics (*e*.*g*., Dice overlap), as global agreement may remain high while local topology is compromised. These limitations motivate a formulation in which coordinate estimation is constrained to the intrinsic surface geometry of the hippocampus.

In the present work, intrinsic coordinates are computed by solving Laplace equations directly on subject-specific hippocampal surface meshes. Native-space surfaces are derived from a laminar representation of hippocampal tissue using an equidistance model implemented in LayNii (Huber et al. 2021), from which a midthickness surface is extracted as a level set representation. Boundary conditions for anterior–posterior and proximal–distal axes are defined using anatomical termini identified during segmentation and assigned to surface vertices. This ensures that both coordinate estimation and boundary conditions are defined intrinsically on the surface manifold.

Laplace equations are then solved on the native surface mesh to obtain smooth harmonic coordinate fields spanning the anterior-posterior and proximal-distal axes. Because the solution is constrained to the surface manifold, coordinate fields follow the intrinsic topology of the hippocampus and cannot traverse adjacent folds through the volume.

This surface-intrinsic formulation eliminates topological shortcuts introduced by volumetric discretization and improves robustness to small segmentation errors that might otherwise create gaps or bridges between adjacent folds. These benefits are particularly important in regions of fine-scale folding where sulcal widths approach voxel resolution limits.

#### Coordinate uniformization

Following estimation of intrinsic coordinates via the surface Laplace–Beltrami formulation, we apply a monotonic reparameterization to improve sampling uniformity in the unfolded domain. In prior implementations, coordinate values reflected the underlying geometry of the hippocampus, resulting in a non-uniform distribution of vertices along the anterior–posterior and proximal–distal axes. In particular, regions such as the medial uncus and posterior tail occupy relatively small spatial extents compared to the hippocampal body, leading to uneven vertex density across the coordinate domain.

To address this, we remap coordinate values to a uniform distribution while preserving their relative ordering. Specifically, coordinate values at non-boundary vertices are sorted and reassigned to evenly spaced values in the interval [0,1][0,1], leaving boundary conditions unchanged. This transformation preserves the topology and ordering of the coordinate system while ensuring a more uniform sampling of vertices in unfolded space.

In prior implementations, non-uniform coordinate distributions were compensated for by using an adaptively spaced mesh in the unfolded domain, designed to produce more uniform vertex spacing after transformation to native space. In the present formulation, uniformization of the coordinate system allows the use of a regular mesh in unfolded space while still achieving uniform sampling in native space. This results in more evenly distributed mesh elements following surface resampling, improving numerical stability and reducing variation in face sizes across the surface.

#### Surface-based correspondence via unfolded-space resampling

Prior HippUnfold versions established correspondence between unfolded and native space through volumetric transformations defined by intrinsic coordinate fields. In this formulation, a canonical unfolded surface was mapped into subject-specific native space using inverse interpolation (SciPy’s griddata (Virtanen et al. 2020)). Because this transformation is defined in the volumetric domain, vertex positions must be interpolated within a discretized 3D deformation field. This interpolation is not constrained to the surface manifold and can introduce geometric distortion, particularly in regions of complex folding.

In the present formulation, correspondence is instead established through surface-based resampling in the unfolded domain. Following estimation of intrinsic coordinates, each subject’s hippocampal surface is parameterized in the unfolded coordinate system. Vertex correspondence across subjects with respect to topology is therefore defined intrinsically in the unfolded domain. An additional refinement stage of unfolded, 2D registration is then performed using the following features: thickness, gyrification, and curvature (DeKraker et al. 2023).

Subject-specific surfaces are brought into correspondence by resampling each native hippocampal surface directly onto a standardized mesh using Connectome Workbench surface resampling (wb_command - surface-resample) (Glasser et al. 2013)). In this operation, the native subject surface is treated as the surface to be resampled, while the subject’s registered unfolded surface and the reference unfolded surface provide the source and target parameterizations, respectively. Vertex positions are interpolated using barycentric interpolation defined by the correspondence of the flat surfaces, yielding a subject-native surface at the standard mesh density with consistent vertex correspondence across individuals. Because interpolation is constrained by the surface parameterization rather than a volumetric deformation field, this approach preserves surface topology and avoids the misplaced or outlier vertices that can arise from inverse volumetric warping. This is analogous to surface resampling used for spherical cortical surfaces in the Human Connectome Project and related pipelines, except that here the common parameterization is a two-dimensional unfolded representation of the hippocampus rather than a sphere.

#### Inner and outer surfaces

In prior implementations, inner and outer hippocampal surfaces were obtained by applying inverse volumetric transformations to canonical surfaces, using the same deformation fields that mapped unfolded coordinates to native space. As with the midthickness surface, this approach relied on interpolation within a volumetric warp field and was therefore susceptible to geometric distortion and inconsistencies in vertex correspondence.

In the present framework, vertex correspondence is defined intrinsically on the midthickness surface, and inner and outer surfaces must be constructed in a manner that preserves this correspondence. One possible approach would be to independently generate inner and outer surfaces and solve intrinsic coordinate systems on each surface, enabling surface-based resampling analogous to the midthickness. However, this would require defining consistent boundary conditions and solving Laplace equations separately for each surface, introducing additional complexity and potential inconsistencies across laminar depths.

Instead, we construct inner and outer surfaces by deforming the native midthickness surface using volumetric registration between laminar boundaries. Specifically, displacement fields derived from the laminar representation are used to map the midthickness surface toward the inner (SRLM) and outer hippocampal boundaries, while preserving vertex correspondence. This approach ensures that all surfaces share a common mesh topology and vertex indexing, enabling consistent sampling across laminar depth.

Because inner and outer surfaces are derived from the midthickness rather than independently parameterized, this approach maintains correspondence but may introduce small deviations from true laminar geometry, particularly in regions of high curvature. As a result, measures such as cortical thickness derived from these surfaces should be interpreted with this approximation in mind. Nevertheless, this strategy provides a practical balance between geometric fidelity and consistent vertex correspondence across surfaces.

### Evaluation of performance

To evaluate the impact of the surface-intrinsic formulation, we compared results obtained using the original HippUnfold framework (volumetric formulation) and the present surface-intrinsic approach. Rather than isolating individual components, we assess the combined effect of these changes on surface geometry and correspondence.

Quantitatively, the two formulations were compared directly on the same participants across multiple sessions and scanners: MICs 3T dataset (n=50, 2 sessions) and PNI 7T dataset (n=10, 3 sessions) (Royer et al. 2022; Cabalo et al. 2025). In the PNI dataset, T1 images were acquired with an MP2RAGE sequence with the following parameters: voxel size 0.5 mm^3^, repetition time 5,170 ms, echo time 2.44 ms, TI1 = 1000 ms, TI2 = 3200 ms, flip angle 4°, iPAT acceleration factor 3, partial Fourier 6/8, 320 sagittal slices, with a final matrix size 320 × 320 × 320. In the MICs dataset, T1 scans were acquired with an MP2RAGE sequence with the following parameters: voxel size 0.8 mm^3^, repetition time 5,000 ms, echo time 2.9 ms, TI1 = 940 ms, T12 = 2830 ms, flip angle 5°, iPAT acceleration factor 3, bandwidth 270 Hz per px, partial Fourier 6/8, 240 sagittal slices, with a final matrix size of 320 × 320 × 240. We then examined consistency, identifiability, and mesh quality of the midthickness surfaces.

Surfaces were linearly transformed to MNI152 space to remove differences in global scale. Vertex coordinates were then zero-centered, and left hippocampi were reflected to match right hemispheres, leaving only diffeomorphic differences between surfaces. Pairwise Pearson correlations were computed between all subject–session surface xyz coordinates. Consistency was defined as the correlation between repeated sessions of the same subject. Identifiability was defined as the difference between within-subject (session–session) and between-subject correlations, normalized by consistency. Mesh quality was assessed using the scaled Jacobian computed with the Python library PyVista, which summarizes geometric properties including face regularity, edge lengths, and the presence of inverted elements. Statistical differences between hemispheres and surface versus volume formulations were assessed using repeated-measures ANOVA.

#### Temporal lobe epilepsy classification

Temporal lobe epilepsy is one of the most common drug-resistant epilepsies in adults, and is associated with hippocampal pathology (Blümcke et al. 2013; Bernasconi et al. 2003; Bernhardt et al. 2015; Cendes et al. 1993). Previous work showed that surface-shape mapping and HippUnfold outputs can help to diagnose and lateralize temporal lobe epilepsy, even in otherwise MRI-negative cases (Bernhardt et al. 2016; Caldairou et al. 2021; Ripart et al. 2024).

To assess whether improvements in surface geometry and correspondence translate to downstream analyses, we compared performance in a classification task: lateralization of temporal lobe epilepsy (TLE). Performance was compared between the volumetric and surface-intrinsic formulations in a cohort of n=65 unilateral pharmaco-resistant TLE patients (mean age 37.5±11.7, 33M/32F, 39 left and 26 right-lateralized).

These patients were recruited and scanned at the Montreal Neurological Institute-Hospital, under the same protocol described above (Royer et al. 2022). Analyses were repeated using logistic regression and a support vector machine using 5-fold cross validation with bootstrapping (n=100) to ensure fair and independent evaluation. Performance was evaluated as the area under the curve of a receiver operating curve (AUC ROC), which weighs both sensitivity and specificity.

#### Robustness to corrupted images

Previous implementations reported a failure rate of approximately 1% for standard T1-weighted images (DeKraker et al., 2022). To further assess robustness, we corrupted sample images by injecting Gaussian noise or blurring to determine the conditions under which performance degrades or fails. These corruptions were implemented using torchio (Pérez-García et al. 2021). Noise amplitude was defined relative to the image’s robust intensity range, computed as the difference between the 2nd and 98th intensity percentiles, with standard deviations set to fixed proportions of this range. Spatial blurring was implemented via a Gaussian kernel with specified voxel-scale standard deviation ranges. Performance was quantified using the consistency metric described above in n=20 samples (10 left and 10 right hippocampi).

## Results

All updated HippUnfold code is available online at https://github.com/khanlab/hippunfold, with details of the updates at https://github.com/khanlab/hippunfold/releases/tag/v2.0.0. Updated Documentation can be found at https://hippunfold.khanlab.ca/en/dev-v2.0.0/.

Manual inspection of updated HippUnfold results was performed by neuroanatomical experts. Healthy control scans across various datasets, including HCP (Van Essen et al. 2012), PNI (Cabalo et al. 2025), and MICs (Royer et al. 2022), were examined by authors JD and AK. Prior implementations reported occasional QC failures (∼1%) in standard T1w datasets (DeKraker et al. 2022), but no failures were observed using the updated software across the datasets examined here.

Qualitative comparison between the prior volumetric (version 1) and present surface-intrinsic formulations (version 2) is shown in **Figure 2A**. The surface-intrinsic formulation demonstrates a lack of misplaced vertices, better centering of the midthickness surface on hippocampal tissue, improved visibility of gyral/sulcal folding, and minimized distortion between folded/unfolded space. Because the unfolded representation is sampled using a regular mesh, face sizes are uniform in unfolded space. Checkerboard visualization further highlights the reduced variation in face scaling between folded and unfolded representations, indicating improved preservation of local geometric relationships during parameterization.

**Figure 2.**
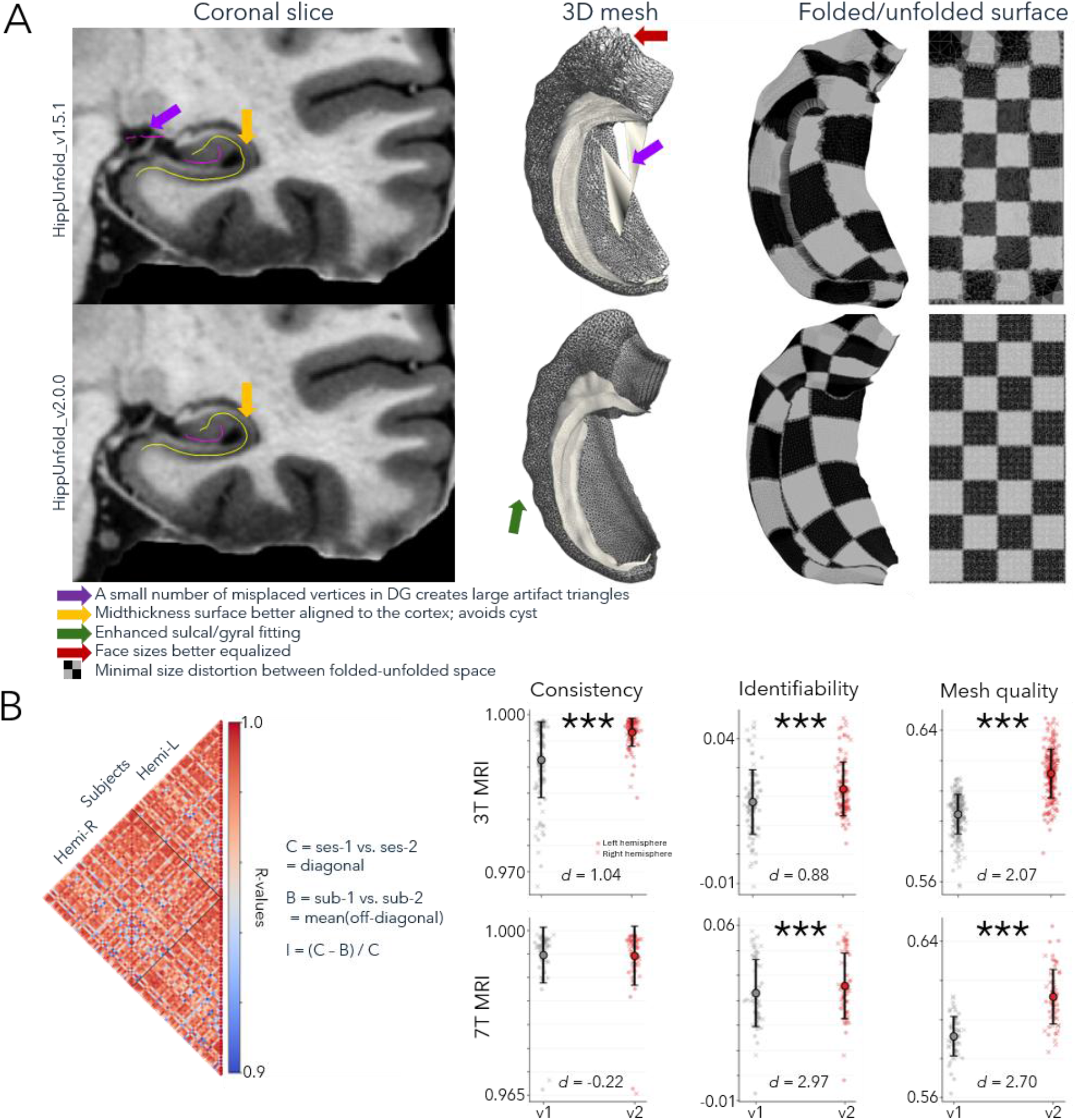
HippUnfold version comparison. **A)** Qualitative comparison between a healthy participant run through HippUnfold version 1 and version 2. Notable features are highlighted by the coloured arrows. **B)** Quantitative comparison between HippUnfold versions with repeated sessions with 3T and 7T MRI data, each with multiple sessions. Derivation of measures is shown to the left, and quantitative results are shown to the right. Circles indicate left hemispheres, x’s indicate right. Red indicates version 2, grey indicates version 1. *d* indicates paired-samples Cohen’s D measure. ^***^ indicates p<0.001.

Quantitative comparisons between the prior volumetric and present surface-intrinsic formulations across 3T and 7T MRI cohorts are shown in **Figure 2B**. Consistency, identifiability, and mesh quality were significantly improved using the surface-intrinsic formulation, except for consistency in the 7T MRI cohort, where no significant difference was observed. This likely reflects a ceiling effect, as both formulations demonstrated near-perfect consistency (R=0.995) in the 7T cohort. Such reproducibility is notable given the substantial session-to-session variability typically introduced by MRI acquisition and downstream automated processing pipelines (Botvinik-Nezer et al. 2020; Glatard et al. 2015). No hemispheric differences were observed in consistency or mesh quality, but a main effect of hemisphere was seen for identifiability in both 3T and 7T datasets (p<0.05).

We next evaluated whether improved surface representation translated to enhanced lateralization in 65 patients with unilateral temporal lobe epilepsy (**Figure 3**). Classification performance improved across nearly all morphometric features, as well as when combining all features, relative to the prior volumetric formulation. The sole exception was surface area, for which the volumetric formulation performed slightly better in logistic regression models only.

**Figure 3.**
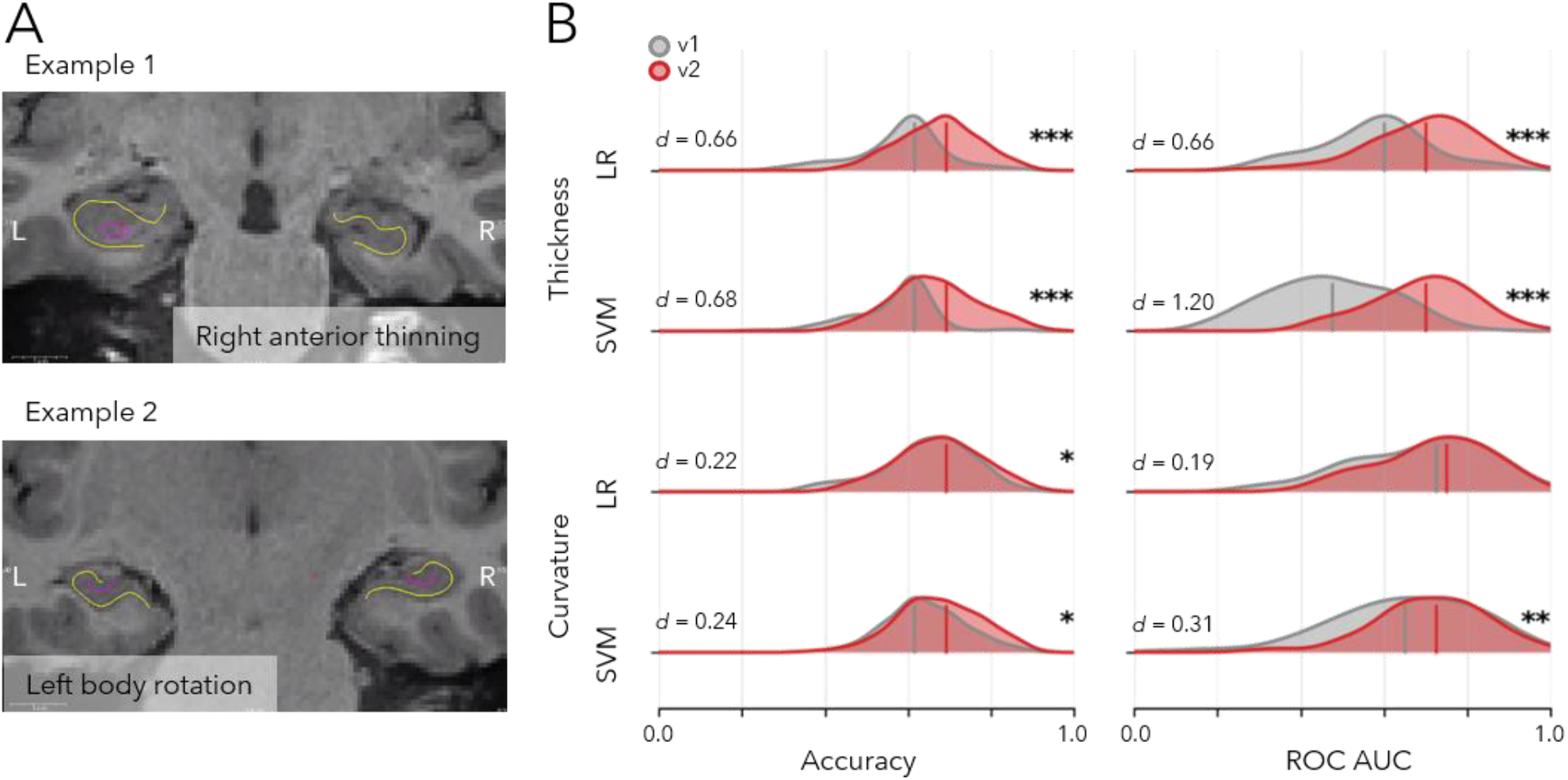
Temporal lobe epilepsy lateralization in HippUnfold version 1 vs. version 2. **A)** Examples of subtle thickness- and curvature-based features for lateralizing TLE. **B)** Logistic regression (LR) and support vector machine (SVM) classification of left vs. right TLE based on thickness and curvature in HippUnfold version 1 (grey) and version 2 (red). ROC AUC stands for receiver operating characteristic area under the curve. *d* indicates paired-samples Cohen’s D measure.

**Figure 4.**
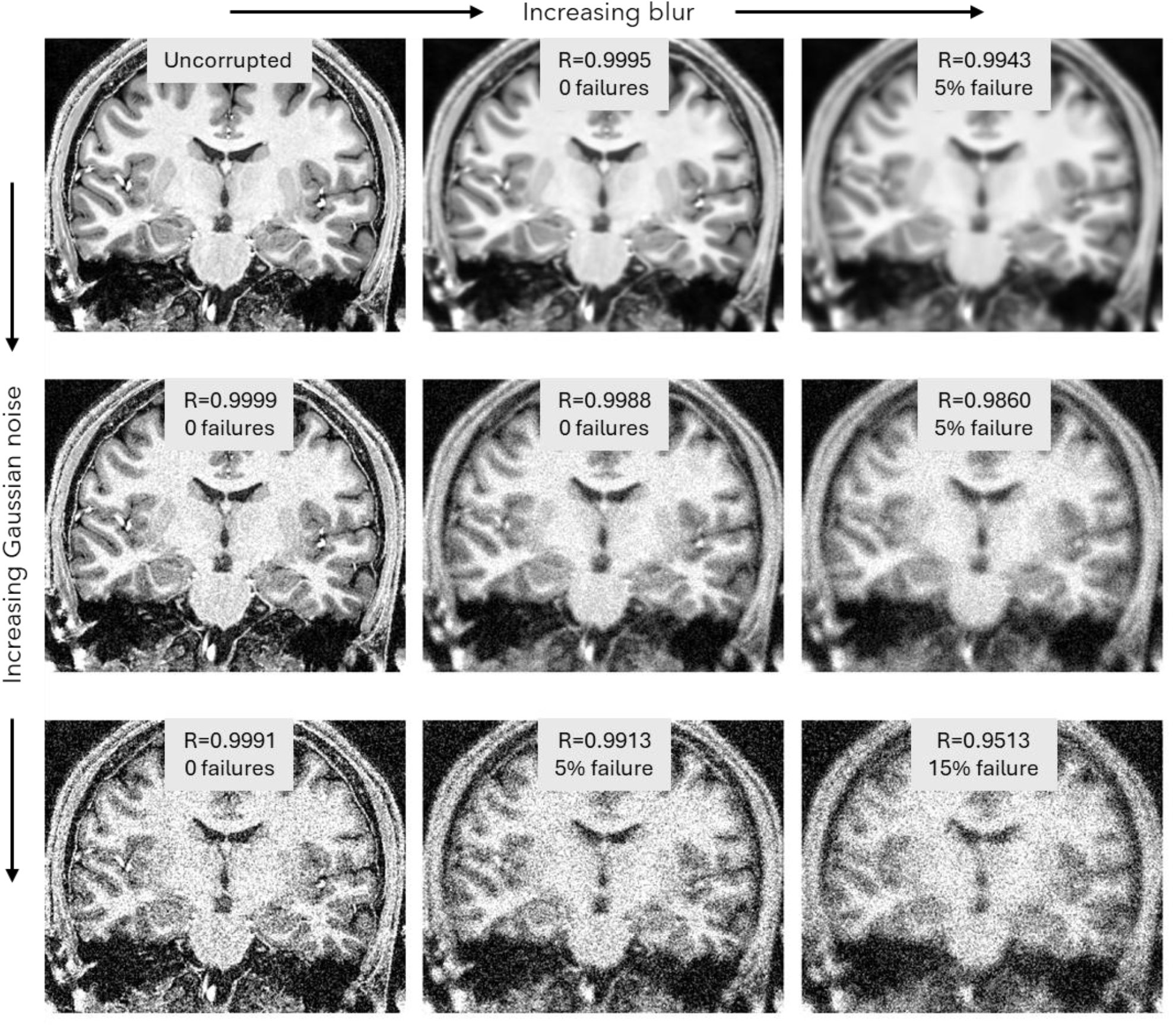
Testing robustness to image corruption. Images were corrupted by applying Gaussian noise at 10% or 20% of the image intensity range, or by a Gaussian blur with a sigma randomly sampled between 0.5-0.8 voxels or 1.0-1.5 voxels. As with consistency measures, performance was evaluated by surface xyz coordinates (n=10L, 10R), in this case relative to each subject’s uncorrupted image.

HippUnfold was originally developed using high quality Human Connectome Project (Van Essen et al. 2012), 7T MRI (DeKraker et al. 2018), and *ex vivo* histological datasets (DeKraker et al. 2023). As a result, a common question from users is whether the framework generalizes to lower-quality MRI acquisitions that are often more practical in specialized populations or challenging imaging environments (Bernasconi et al. 2019; Aklamanu 2025). To evaluate robustness under such conditions, we simulated image degradation by systematically corrupting input images and assessing the point at which performance began to deteriorate.

## Discussion

In the present work, we reformulated hippocampal unfolding as a surface-intrinsic problem in which geometry, intrinsic coordinates, and inter-subject correspondence are defined directly on the hippocampal surface manifold. This reformulation substantially improves surface consistency, identifiability, and mesh quality across both 3T and 7T MRI datasets, while also improving downstream performance in a clinically relevant temporal lobe epilepsy lateralization task. Collectively, these findings demonstrate that defining hippocampal unfolding directly on the surface manifold improves both the anatomical fidelity and practical utility of hippocampal surface representations.

A major challenge in hippocampal segmentation is establishing anatomical validity. Most existing approaches rely on manual segmentation protocols derived from histological reference data, including harmonization efforts such as the Hippocampal Subfields Group (HSG), which aim to develop consensus heuristics that can be applied reproducibly by expert raters on MRI (Olsen et al., 2019; Yushkevich et al., 2015). Automated tools such as ASHS and related atlas-based methods similarly depend on training from these manually delineated labels (Yushkevich et al. 2016; Wisse et al. 2016). In contrast, the HippUnfold framework takes a fundamentally different approach: rather than translating histological anatomy into simplified heuristic segmentation rules, we seek to preserve anatomical validity by directly leveraging *ex vivo* and histological ground truth data and improving the anatomical correspondence between hippocampi such that these data can be mapped faithfully across individuals. The present surface-intrinsic reformulation strengthens this philosophy by improving the fidelity with which subject-specific folding patterns and local topology are preserved during alignment, thereby providing a more anatomically principled basis for transferring ground-truth anatomical information across subjects.

These improvements may be particularly valuable for initiatives such as HippoMaps (DeKraker et al. 2025), which aim to integrate hippocampal data across modalities and studies within a shared surface framework, and provide a promising avenue for aligning results between labs, saving costly data acquisitions, and enabling discovery of new relationships between modalities. More anatomically faithful surface representations should enable sharper and more spatially precise multimodal maps in future applications. At the same time, we note that the *ex vivo* and histological datasets incorporated into HippoMaps were generated through iterative refinement of carefully curated manual segmentations followed by surface fitting, to ensure optimal unfolding. Thus, gains from the present automated improvements may be most pronounced in typical *in vivo* MRI applications where segmentation and unfolding errors are more common. To maintain compatibility with prior datasets, HippoMaps has been updated to support interpolation between legacy and current HippUnfold surface representations.

HippUnfold also shows strong promise in clinical applications due to its sensitivity to subtle inter-individual differences. This is particularly relevant in diseases that impact the hippocampus, including schizophrenia (Nelson et al. 1997; Haukvik et al. 2018), depression (Videbech and Ravnkilde 2004; Bremner et al. 2000), epilepsy (Jack 1994; Ripart et al. 2024; Blümcke et al. 2013; Bernhardt et al. 2016; Cendes et al. 1993; Bernasconi et al. 2003), and dementia (Schuff et al. 2009; Convit et al. 1997; Shi et al. 2009). In temporal lobe epilepsy specifically, prior work demonstrated that HippUnfold-derived morphometric features enable highly sensitive lateralization of epileptogenic lesions, including in MRI-negative cases (Ripart et al. 2024), complementing earlier work (Caldairou et al. 2021). Here, we show that improving anatomical fidelity through the surface-intrinsic formulation further enhances this performance. This finding is particularly important for the detection of subtle or early-stage abnormalities, where structural deviations may be highly localized and easily obscured by noise, partial volume effects, or misalignment in conventional volumetric approaches. By more accurately preserving the intrinsic geometry of the hippocampal surface, our method enables finer-grained characterization of morphological features that are likely to be affected in the earliest phases of disease. Earlier studies employing the HippUnfold version 1 could benefit from higher precision if reanalyzed using the present methods. This increased sensitivity may, in turn, support earlier diagnosis, improve tracking of disease progression, and provide a more precise substrate for linking structural alterations to underlying biological mechanisms.

Several opportunities remain for future methodological development. First, while the present work improves the geometric formulation of hippocampal unfolding, the overall pipeline remains dependent on machine learning models for the initial tissue segmentation stage. As with most supervised segmentation approaches, generalization to new datasets depends strongly on whether the imaging domain is sufficiently represented in the training data, including factors such as contrast, resolution, subject population, and pathology. To address this, multiple specialized HippUnfold models have been developed beyond the original HCP young-adult model, including neonatal models (Nichols et al., 2025), while additional models targeting older adult and neurodegenerative cohorts are currently in development. Other promising approaches seek to improve generalizability through contrast-agnostic synthesized training data, as demonstrated by SynthSeg-like methods, an approach that is also being actively investigated within the HippUnfold framework.

Beyond segmentation, several geometric limitations remain in the unfolding formulation itself. While the present work defines intrinsic coordinates and correspondence directly on the midthickness surface, inner and outer laminar surfaces are still derived through deformation of the midthickness rather than independently parameterized. Future work may extend surface-intrinsic principles to these additional surfaces to further improve laminar accuracy and thickness estimation. In addition, the present Laplace formulation still requires predefined anatomical boundary conditions and therefore remains partially dependent on anatomical priors, unlike emerging fully geometry-intrinsic approaches (e.g., Diers et al., 2023). Finally, anterior–posterior and proximal–distal coordinate fields are solved independently, without explicit constraints enforcing orthogonality, representing another potential avenue for refinement in future implementations.

Overall, the present work establishes a more anatomically faithful framework for hippocampal unfolding by defining core components of the pipeline directly on the hippocampal surface manifold. By improving topological fidelity, geometric accuracy, and inter-subject correspondence, this surface-intrinsic formulation enhances the robustness and interpretability of hippocampal surface analysis and provides a foundation for future surface-based neuroimaging applications.

## Acknowledgements

J.D. was supported by a Natural Science and Engineering Research Council of Canada Post Doctoral Fellowship award (NSERC-PDF), a Molson Engineering fellowship of the Montreal Neurological Institute, Helmholtz International BigBrain Analytics and Learning Laboratory (HIBALL), Healthy Brains and Healthy Lives (HBHL), and the Centre for Aging and Brain Health Innovation (CABHI).

B.C.B. was supported by the Canadian Institutes of Health Research (CIHR), NSERC, BrainCanada, HBHL, HIBALL, CABHI, a McGill-Western Initiative for Translational Neuroscience (ITN) grant, and the Centre for Excellence in Epilepsy at the Neuro (CEEN).

A.N. was supported by the Canadian Institutes of Health Research (CIHR).

J.R. is supported by a Banting Postdoctoral Fellowship from NSERC.

B.G.K is supported by a Natural Science and Engineering Research Council of Canada Postdoctoral Research Award (NSERC-CPRA).

D.B. was supported by the fund from the McGill and Western Initiative for Translational Neuroscience (ITN), “Translating epilepsy neuroimaging biomarkers into the operating room”.

M.T.A. and J.T. were supported by grants 346934 (PRIMAL), and 358944 (Flagship of Advanced Mathematics for Sensing Imaging and Modeling) from the Research Council of Finland and a grant from the Jane and Aatos Erkko foundation.

M.F.G. was supported by NIH NIA R01AG092088.

J.C.L. was supported by an NSERC Discovery Grant RGPIN-2023-05562, research start-up funding from the Department of Clinical Neurological Sciences at Western University, and a McGill-Western Initiative for Translational Neuroscience (ITN) grant, a joint funding grant between HBHL and BrainsCAN.

## Conflicts of Interest

J.D. and B.C.B. are co-founders of BrainScores Inc. and hold stock.

